# Electrophysiological signatures of temporal context in the bisection task

**DOI:** 10.1101/2023.03.15.532795

**Authors:** Cemre Baykan, Xiuna Zhu, Artyom Zinchenko, Hermann J. Müller, Zhuanghua Shi

## Abstract

Despite relatively accurate time judgment, subjective time is susceptible to various contexts, such as sample spacing and frequency. Several electroencephalographic (EEG) components have been linked to timing, including the contingent negative variation (CNV), offset P2, and late positive component of timing (LPCt). However, the specific role of these components in the contextual modulation of perceived time remains unclear. In this study, we conducted two temporal bisection experiments, where participants had to judge if a test duration was close to a short or long standard. Unbeknownst to participants, the sample spacing (Experiment 1) and frequency (Experiment 2) were altered to create short and long contexts while keeping the test range and standards the same in different sessions. The results showed that the bisection threshold shifted toward the ensemble mean and that CNV and LPCt were sensitive to context modulation. Compared to the long context, the CNV climbing rate increased in the short context, and the amplitude and latency of the LPCt were reduced. These findings suggest the CNV represents an expectancy wave for upcoming decision-making, while LPCt reflects the decision-making process, both influenced by the temporal context.

## Introduction

Processing the vast amount of information that surrounds us can be challenging, as our sensory organs have limited processing capacity (Wolfe, 1994), and more so, our memory and attentional resources (Cavanagh & Alvarez, 2005). To tackle limitations, our brain has developed ensemble perception (Whitney & Yamanashi Leib, 2018), a method for quickly understanding the essence of a scene by extracting statistical information, such as the mean and variance, from its features. For example, we can quickly determine the average size of apples in a supermarket by just glancing at them, without a need to process each individual apple in detail. We use similar forms of ensemble perception to process basic features, such as the average motion, orientations, and colors (Albrecht et al., 2012; Ariely, 2001; de Gardelle & Summerfield, 2011; Parkes et al., 2001; Piazza et al., 2013; Williams & Sekuler, 1984), as well as sequential durations (Zhu et al., 2021). There are two types of ensemble representations: spatial (Whitney & Yamanashi Leib, 2018) and temporal (Jones & McAuley, 2005; Schweickert et al., 2014). Spatial ensemble representation involves a group of similar objects that are presented simultaneously, while temporal ensemble representation involves processing a sequence of stimuli over time.

When serving as context, both types of ensemble statistics can influence judgments of individual items (Ariely, 2001; Zhu et al., 2021). For instance, a temporal bisection task to judge if a probe duration is close to a fixed ‘short’ or ‘long’ anchor is thought to rely only on the probe’s distance to the standards. However, studies have shown that sampling durations can affect our judgment of related durations (Allan, 2002; Penney & Cheng, 2018; Wearden & Ferrara, 1995). The transition between the short and long durations, known as the bisection point, is often influenced by the mean of the sample durations (Zhu et al., 2021). This form of bias, known as *the spacing effect*, occurs when distances among sampled durations are uneven (Wearden & Ferrara, 1995), and *the range effect*, occurs when the spread of the sample set influences its individual durations (Droit-Volet & Wearden, 2001; Penney et al., 2014; Wearden & Ferrara, 1996). While the behavioral effects of the temporal contextual modulation are now better understood (Zhu et al., 2021), the neural mechanisms governing common temporal context effects are not yet fully understood, although some recent research has shed light on this topic (Damsma et al., 2021; Wiener et al., 2018; Wiener & Thompson, 2015).

Recent EEG studies have identified several event-related potentials (ERP) associated with time processing and contextual modulation (Lindbergh & Kieffaber, 2013; Ng et al., 2011; Wiener & Thompson, 2015). For example, in a bisection task, the contingent negative variation (CNV) - a negative polarity waveform that appears in the supplementary motor area (SMA) - has been found to increase in negativity as the interval progresses and levels off when the duration exceeds the geometric mean of the short and long anchors (Ng et al., 2011; van Rijn et al., 2011; Wiener & Thompson, 2015). Additionally, post-interval positivity ERPs, which appear in the range between 200 to 600 ms after the stimulus offset in the same electrode clusters as CNV after the stimulus offset, have been found to vary with duration judgments and temporal decisions (Damsma et al., 2021; Ofir & Landau, 2022). The early positivity component P2, peaking around 200 ms after the stimulus, has been suggested to be linked with perceived duration length (Kononowicz & van Rijn, 2011), although the relationship between P2 and the probe duration remains unclear (Kononowicz & van Rijn, 2014; Lindbergh & Kieffaber, 2013). The late positivity components, P3, P3b, or late positive component related to timing (LPCt), measured around 300 - 600 ms after the stimulus, have been related to temporal decision-making (Bannier et al., 2019; Lindbergh & Kieffaber, 2013; Paul et al., 2011). For example, Ofir and Landau (2022) revealed that the offset-evoked P3 was negatively correlated with the stimulus duration in a bisection task, and its amplitude decreased as the stimulus duration increased. Using a drift-diffusion model, Ofir and Landau (2022) could predict behavioral performance by assuming that the amplitude of the offset-P3 reflects the proximity of the temporal accumulation to the decision boundary (i.e., the bisection point). Likewise, LPCt has also been found to vary with the probe duration, with larger positive amplitudes being associated with shorter durations (Wiener & Thompson, 2015). Given that LPCt or P3 is measured after the duration offset, higher peaks for the short compared to long intervals are interpreted as a sign that decisions for short intervals are more demanding, as the decision process remains active and unresolved at the offset of a short presentation (Lindbergh & Kieffaber, 2013).

Early studies primarily focused on those ERP components on temporal accumulation (e.g., Macar et al., 1999), which did not take contextual modulation into account. More recent studies have shown those ERP components are also sensitive to temporal contexts. Wiener and Thompson (2015) have shown that both the CNV and LPCt covaried with the duration presented in the prior trial. Damsma et al. (2021) asked participants to reproduce intervals drawn from two different but overlapping ranges (the short and the long). When the same interval was reproduced in the short-range compared to the long-range session, the reproduction elicited higher amplitudes of CNV and offset P2. Furthermore, the amplitudes of the CNV and the offset P2 decreased as the prior interval increased. By probing a bisection task in sub-second and supra-second ranges separately, Ofir and Landau (2022) found that the offset P3 amplitude resembled a similar pattern in the two ranges, highlighting the nature of contextual modulation (Baykan & Shi, 2022).

Decisions in a bisection task can be made *during* stimulus presentation without waiting until the end, as it becomes clear whether a duration is short or long once the elapsed time passes the bisection point between the short and the long. But a decision in a reproduction task can only be made during the late reproduction phase, after the full presentation of the duration. As a result, ERP components related to different timing tasks - bisection or reproduction - may reflect different temporal cognitive processes depending on the task being applied (Gontier et al., 2009; Kononowicz & van Rijn, 2014; van Rijn et al., 2011).

It is important to note that the aforementioned EEG studies primarily focused on neural activities during the probe itself. While some studies have discussed contextual modulation (e.g., Ofir & Landau, 2022), the contexts are often vastly different (e.g., sampling from different ranges of durations, such as subseconds vs. super-seconds). None of these studies have examined *ensemble contexts* within the same range of durations, but with different duration spacing or sample frequencies. Neurophysiological mechanisms underlying such *ensemble contexts* are not yet fully understood. To address this issue, we designed two experiments using the bisection task with manipulations of sampled durations. In both experiments, the short standard was 400 ms, the long standard 1600 ms. Participants had to judge whether a probe duration was close to the short standard or the long standard. Unbeknown to participants, the sampled durations in Experiment 1 were positively skewed in one session and negatively skewed in the other, while in Experiment 2 there were high frequencies of short samples in one session and long samples in the other. Based on previous findings (Ng et al., 2011; Wiener & Thompson, 2015), we expected the CNV peak latencies would correlate to the internal decision criterion of the bisection task, where peak latencies would be earlier in short compared to long contexts, but would plateau after the ensemble mean duration. According to literature (Kononowicz & van Rijn, 2014; Tarantino et al., 2010), which suggests that the P2 amplitude is linked to the stimulus magnitude, we hypothesized that the P2 amplitude would increase with target interval increases and be more positive in short relative to long contexts. To distinguish the late positivity components, e.g. LPCt, from the P2 component, we inserted a 300 ms blank interval after the stimulus offset in Experiment 2, examining the relationship between the LPCt amplitude and the target interval.

### Experiment 1

In Experiment 1, we manipulated the temporal context using the positively skewed (PS, more short durations) and negatively skewed (NS, more long durations) sample distributions, based on our previous work (Zhu et al., 2021). Behaviorally, we expected the same outcome as the previous study - intermediate durations would be more likely to be judged as “long” in the PS than the NS context.

## Method

### Participants

20 participants with no hearing impairment took part in Experiment 1 in exchange for a monetary reward or course credit at LMU Munich. The sample size was calculated based on the effect size of a similar temporal bisection study (Zhu et al., 2021) with η_*g*_ = .26, and the assumption of α = .05 and power 1 − β = .95, which required a sample size of 16 participants. To be safe for EEG analysis, we increased the sample size to 20. All participants provided written informed consent before their participation. One participant was excluded in the formal analysis because of excessive eye and body movement artifacts. Thus, the final sample in Experiment included 19 participants (10 females, mean age 27.2 years, *SD* = 4.2 years), who were naive to the purpose of the study. The study was approved by the Ethics Board of the Department of Psychology at LMU Munich.

### Stimuli and Procedure

The auditory stimuli were generated using the PsychoProtAudio library and presented through loudspeakers (Logitech Z130) using the Psychtoolbox 3 (Kleiner et al., 2007). Instructions and feedback text were displayed on a CRT monitor.

Participants sat in a sound-attenuated, moderately lit test room. Prior to the formal experiment, participants received a practice block consisting of 5 presentations of the short and long standards (400 and 1600 ms) to familiarize themselves with the standards. During the practice, participants made “short” or “long” judgments and received feedback on whether they were correct or incorrect. In the formal test, each trial started with a visual fixation and a brief beep (20 ms, 1000 Hz, 60 dB), followed by a 500 ms blank display, signaling the start of a new trial. A white-noise stimulus (60 dB) was then presented for a given duration chosen from the experimental stimulus sets (see below). Immediately after the sound presentation, a question mark appeared, prompting participants to respond by pressing the right or left arrow keys on the keyboard using two index fingers, indicating if the presented sound was close to the short or the long, respectively (Fig. 1a).

**Figure 1.**
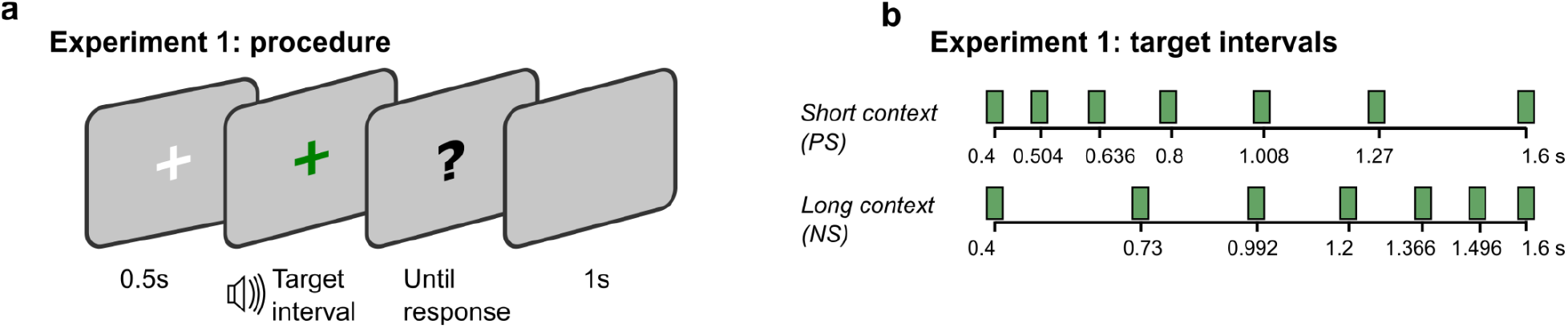
**a)** Each trial started with fixation-cross for 500 ms. It was followed by a target interval presentation. Right after the presentation, a question mark appeared, prompting participants to respond. The inter-trial-interval was 1000 ms. **b)** The target intervals used in Experiment 1. In the short context session (PS), intervals were logarithmically spaced between 400 ms and 1600 ms and the intervals were mirrored in the long context session (NS). Each target interval was presented 48 times during the session.

There were two sessions with each session of 336 trials (six blocks of 56 trials each). Two sessions had different duration sets: the positively skewed (PS) duration set consists of [400, 504, 636, 800, 1008, 1270, 1600] ms, the negatively skewed (NS) duration set [400, 730, 992, 1200, 1366, 1496, 1600] ms (Fig. 1b). The ensemble mean of the NS was 223 ms longer than the ensemble mean of the PS context. Each duration was randomly tested 48 times. The order of sessions was counterbalanced across participants (before the outlier exclusion).

### EEG acquisition and analysis methods

Electrical brain activity was recorded from 64 scalp locations (actiCAP system; Brain Products, Munich, Germany) using the BrainVision Recorder software (Brain Products GmbH, Munich, Germany) and a BrainAmp amplifier (DC to 250 Hz) at the sampling rate of 1000 Hz. During the experiment, the impedances of all electrodes were kept below 10 kΩ. The electrode FCz was used as an online reference and was re-referencing to temporal-parietal electrodes offline (TP9 and TP10).

The EEG data were analyzed using BrainVisionAnalyzer 2.0 software, with a bandpass filter of 0.1 to 70 Hz. Artifacts caused by eye blinks, eye movements, and muscle noises were removed using independent component analysis (ICA) and visual identification. Before segmentation, the continuous EEG data were inspected automatically using the raw data inspection procedure in the analyzer software and were bandpass-filtered from 0.1 to 30 Hz.

### ERP components

All ERP components reported here were calculated for each participant, target interval, and temporal context. The onset-locked ERP data for CNV analyses were baselined to the average voltage 200 ms prior to the stimulus onset, using six clustered frontocentral electrodes FCz, FC1, FC2, C1, C2 and Cz (Kononowicz & van Rijn, 2014; Ng et al., 2011). To examine the evolving velocity of the CNV negativity over time, we conducted linear regression in the time window from the end of post-onset P2 (250 ms) to the start of the CNV (800 ms) as per previous literature (Ng et al., 2011) and obtained slopes for individual participants in each condition. We extracted the CNV peak latencies as the minimum (most negative) amplitude from stimuli onsets to the longest duration offset (i.e., 1600 ms) for each target interval in each context. We defined the mean CNV amplitude of each target interval as the average waveform in the interval starting from the late negativity onset (250 ms after the onset) and having a length of the stimulus duration (Kruijne et al., 2021). The stimulus offset P2s were calculated using the same frontocentral electrodes as used for the CNV analysis (Damsma et al., 2021). The offset-locked ERP data for P2 analyses were baselined to the 100 ms time window surrounding the stimulus offset (50 ms preceding and 50 ms following the offset) (Kononowicz & van Rijn, 2014). We extracted the P2 peak latencies as the maximum (most positive) amplitude within the 0 - 500 ms duration offset window. We defined the mean P2 amplitude of each target interval as the average waveform between 140 and 300 ms after the stimulus offset (Kononowicz & van Rijn, 2014).

### Data analysis

Psychometric functions were estimated using the logistic function with the Quickpsy package in R (Linares & López-Moliner, 2016), the point of subjective equality (PSE) was then calculated at the threshold of 50%, and the just noticeable difference (JND) as the difference between the thresholds at 50% and 75%. Mean PSEs and JNDs were compared using repeated-measures analysis of variance (ANOVA), if necessary, with additional Bayes factor *(BF*) analyses to provide a more rigorous assessment of the null hypothesis (Rouder et al., 2009). For analysis of the EEG components, we applied a linear mixed model, which can accommodate the covariant factor (duration) in addition to the fixed effects addressed by ANOVA. Mixed models are robust to violations of sphericity and do not inflate Type I errors (Singmann & Kellen, 2019). The *p*-values reported for mixed models were calculated using the Kenward-Roger approximation.

## Results

### Behavioral results

Figure 2 illustrates the averaged psychometric functions, mean PSEs, and JNDs. The mean PSE (± standard error, *SE)* for the short-context PS session was significantly shorter (888.7 ± 45.9 ms) than the long-context NS session (958.9 ± 46 ms), *F(*1, 18) = 6.90, *p* = .017, 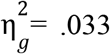, *BF* = 3.04 (see also Table 1). In other words, the same duration (e.g., 1 sec) was perceived as longer in the short relative to the long context. The sensitivities of the bisection, measured by JNDs, were comparable between two sessions, *F*(1, 18) = 2.44, *p* = .14, 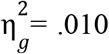, *BF* = 0.74, *g* indicating the spacing of the target intervals did not change the discrimination sensitivity. Thus, The behavioral results are in line with the previous findings (Zhu et al., 2021).

**Table 1.** Mean ERPs and Behavioral PSEs and JNDs

| Experiment | Context | CNV |  |  |  | Behavioral |  |
| --- | --- | --- | --- | --- | --- | --- | --- |
| | | Climbing Rate<br>($\mu\text{V/s}$) | Peak latency<br>(ms) | Peak amplitude<br>($\mu\text{V}$) | Mean amplitude<br>($\mu\text{V}$) | PSE<br>(ms) | JND<br>(ms) |
| 1: Spacing | Short (PS) | **-23** | **773.2** | -6.1 | -1.47 | **888.7** | 121.1 |
|  | Long (NS) | **-20** | **875.9** | -5.6 | -1.17 | **958.9** | 131.7 |
| 2: Frequency | Short (DF) | **-19** | 942.8 | **-5.4** | -0.78 | **749.3** | **96.3** |
|  | Long (AF) | **-17** | 876.6 | **-4.5** | -0.27 | **951.2** | **121.1** |

*Note.* The data are grouped by temporal contexts in both Experiments 1 and 2. Bold values indicate a significant difference between the two contexts.

**Figure 2.**
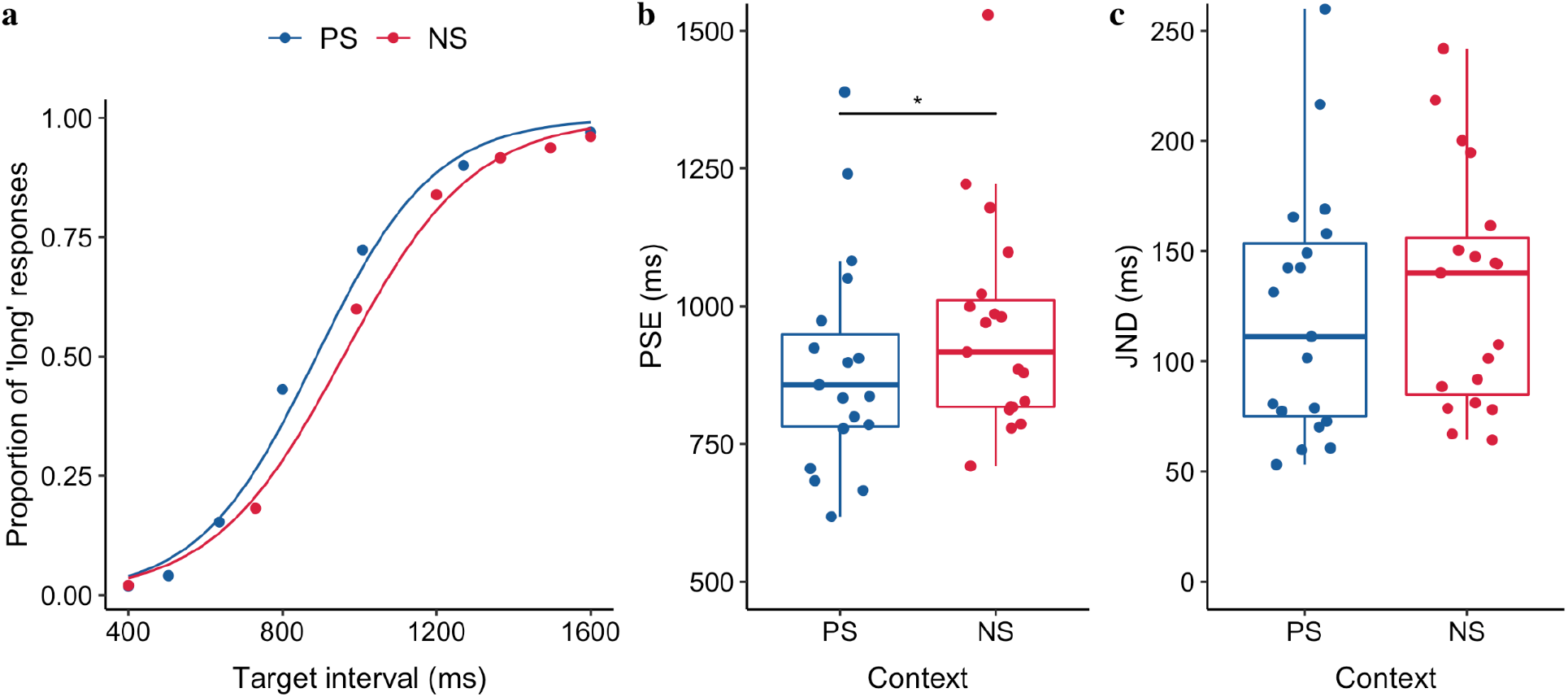
(**a)** The averaged proportion of ‘long’ responses (scatter dots) and the fitted psychometric curves over 19 participants, separated for the positively (PS) and negatively skewed (NS), stimulus-spacing conditions. (**b)** Boxplots of the points of subjective equality (PSEs) of the duration judgments for the PS and NS sessions (* p <. 05). The dots depict individual PSEs. The lower and upper tips of the vertical lines correspond to the minimum and maximum values, the box the interquartile range (between 25% and 75%), and the horizontal line the median. (**c)** Boxplots of JND of the duration judgments for the PS and NS sessions. The dots depict individual JNDs.

### Electrophysiological Results

#### Contingent negative variation (CNV)

Figure 3 illustrates the CNV activities in the short PS (a) and long NS contexts (d), showing the negativity changes over time for different target intervals. To characterize the CNV components, we looked into its formation rate, peak latency, amplitude, and the mean latency. Given the negative ballistic deflation of the activities after P2, we used linear regression to estimate the rate (i.e., slope) at which the CNV was forming within the time window from 250 (after P2) to 650 ms (the start of the CNV). The mean values are listed in Table 1.

**Figure 3.**
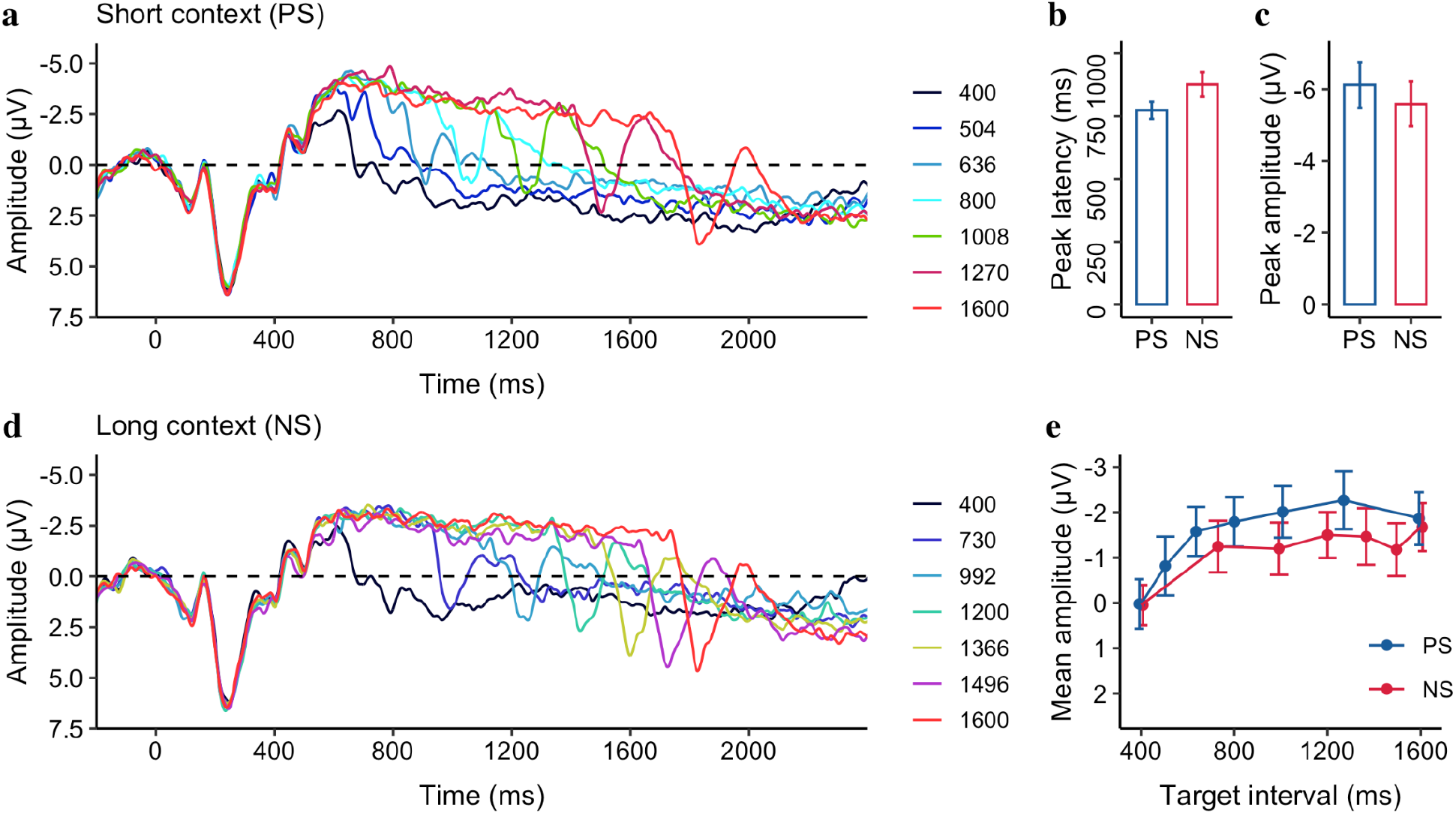
Grand average of the ERP waveforms over the medial frontal electrodes (FCz, FC1, FC2, C1, C2, and Cz), separatedfor different target intervals, separated for (**a**) the short (PS) and (**d**) the long (NS) contexts. The mean CNV peak latency (**b**) and amplitude (**c**) of the target intervals, separated for the PS and NS conditions. (**e**) The mean CNV amplitude as a function of the target interval, separated for the PS and NS conditions. Error bars represent the standard error of the mean.

We found the rate was significantly negative for both the PS context (−23 ± 1.9 μV/s, 95% CI = [−27 to −19] μV/s, *t* (18) = −12.24, *p* < .001) and the NS context (−20 ± 1.6 μV/s, 95% CI = [−24 to −17] μV/s, *t* (18) = −13.12, *p* < .001), but significantly smaller in the PS compared 2 to the NS, *F*(1, 18) = 9.03, *p* = .01, 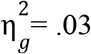, *BF* = 5.53, indicating a faster temporal accumulation in general for durations in the PS relative to the NS session, consistent with the prior research (Macar & Vidal, 2004). Moreover, the CNV peaked significantly earlier for the short context PS relative to the long context NS (773.2 ms vs. 875.9 ms, *F(*1, 18) = 4.99, *p* = 0.04, 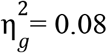, *BF* = *g* 2.17), but only numerically higher in amplitude for the PS relative to the NS (−6.1 μV vs. −5.6 μV, 2 *F*(1, 18) = 3.61,*p* = 0.07, 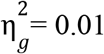, *BF* = 1.12).

As research has shown the mean amplitude of CNV to be correlated with the sample duration (Macar et al., 1999; Pfeuty et al., 2003, 2005), we estimated the mean amplitude separately for individual durations, as depicted in Fig. 3e. We then applied a linear mixed model to the mean CNV amplitude, with the Context as the fixed effect and Duration a covariant effect. The mixed model showed that the mean negativity increased by 1.23 μF per second of Duration *(b* = −1.23, *CI* = [−1.72, −0.73], *p* < .001). However, there was no significant difference between the short and long contexts *(p* = .53) and no significant interaction between the Duration and Context *(p* = .31).

#### Offset P2

Figures 4a and 4c depict the ERP waveforms over the medial frontal electrodes relative to the offset of the stimuli for the short and long contexts, showing a positive peak around 200 ms after the stimuli offset that correlates with the target interval. Figures 4b and 4d show the peak latency and mean amplitude of the offset P2 as a linear trend of the target interval, separated for the PS and NS contexts: The latency decreases, but the amplitude increases as the target interval increases, while there was not much difference between the two contexts. A linear mixed model with the Context as the fixed effect and Duration as a covariant effect was applied to the offset P2 peak latency, which revealed that the peak latency decreased by 78 ms/s of Duration *(CI* = [−144.92, −10.38], *p* = .026). But the peak latency showed no significant difference between the PS and NS Context *(p* = .28). Moreover, there was no significant interaction between Duration and Context *(p* = .58). Similar linear mixed model applied to the Offset P2 mean amplitudes (Fig. 4d) revealed a significant main effect of Duration (*b* = 2.90, *CI* = [1.98, 3.81], *p* < .001). Again, there was no significant Context *(p* = .28) and no interaction between the Duration and Context *(p* = .41). The findings indicate that while the offset P2 was responsive to the target interval, it was insensitive to variation of ensemble contexts.

**Figure 4.**
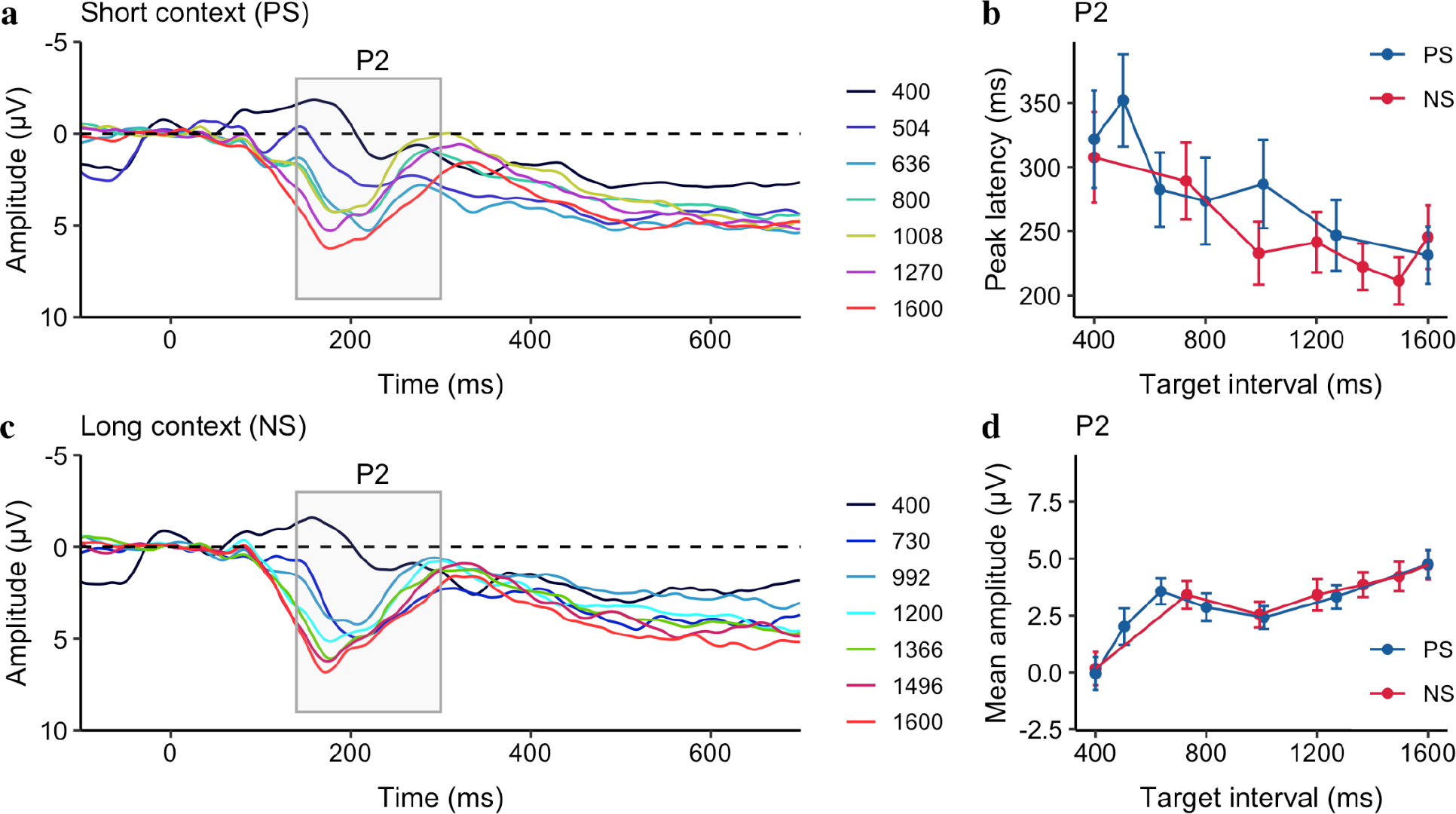
Grand average of the ERP waveforms over the medial frontal electrodes (FCz, FC1, FC2, C1, C2, and Cz), separatedfor different target intervals, separatedfor the PS (**a**) and the NS conditions (**c**). (**b**) Mean P2 peak latency and (**d**) mean P2 amplitude as a function of the target interval, separated for the PS (blue) and NS (red) contexts. Error bars represent the standard error of the correspondent mean.

## Discussion

Experiment 1 replicated previous research (Wearden & Ferrara, 1995; Zhu et al., 2021), confirming that temporal bisection is subjective to the target spacing. Intervals in the short context (PS), relative to the long context (NS), tended to be judged longer, indicating that participants not merely compared the probe duration to the short or long standards (which were the same in both contexts), but also took into account the spacing of the sample durations.

Experiment 1 revealed that the mean amplitude of CNV was linked to the target duration. As the duration increased, so did the mean amplitude. Figure 3e also shows that the mean amplitude leveled off at middle durations (800 to 1200 ms), which is in line with previous research (Macar & Vidal, 2003; Ng et al., 2011) that found CNV plateaued at the geometric mean of the short and long intervals in the bisection task. The findings support that CNV stands for temporal expectation (Amit et al., 2019; Praamstra et al., 2006). Some researchers have suggested that the CNV amplitude is subjective to the context. For example, adapting to shorter durations would lead to an increase in the amplitude of CNV, while adapting to longer durations would decrease the amplitude of the CNV (Li et al., 2017). Here we found the mean (or peak) amplitude of the CNV was higher for the short context (PS) than for the long context (NS), but the difference did not reach statistical significance. On the other hand, we did find that the latency of the CNV was earlier for the short context relative to the long context, which aligns with previous research showing a faster development of the CNV activity for short than long target durations (Pfeuty et al., 2005). However, as Kononowicz and Penney (2016) have suggested, timing isn’t the only factor contributing to the CNV. More complex processes, such as preparation for an upcoming event, could play a role. Therefore, in some cases, the CNV may not truly reflect the temporal interval itself, as revealed in a previous study that CNV-like negativity simply disappears for intervals longer than 4 seconds (Elbert et al., 1991).

Additionally, the offset P2, a common component associated with temporal accumulation as reported previously (Kononowicz & van Rijn, 2014; Tarantino et al., 2010), had a negative correlation with the target interval, which is consistent with previous findings (Kononowicz & van Rijn, 2014). This can be explained by the predictive coding account (Friston & Kiebel, 2009; Kononowicz & van Rijn, 2014; Rao & Ballard, 1999), because short intervals that stopped before the decision threshold (i.e., the bisection point) led to larger ‘prediction errors’ than long intervals, resulting in early P2 latencies. This is also in line with the previous studies showing that short durations lead to longer reaction times (e.g., Bannier et al., 2019). However, the offset P2 was not affected by the spacing modulation, which is in contrast to previous reports that the late positive component of timing (LCPt), peaking at around 300 ms post-offset (later than P2), can be influenced by the task difficulty (Paul et al., 2011), the prior trial duration (Wiener & Thompson, 2015), or the sample set (Ofir & Landau, 2022).

It’s worth noting that previous studies that used bisection or duration comparison tasks often employed durations longer than 800 ms (e.g., Ng et al., 2011), meaning that the expectation of a binary decision would not occur earlier than 500 ms, at which point the CNV is just emerging (as seen in Figure 3). Here we used two short intervals (400 ms, 504 ms), which caused the offset of CNV to happen earlier for preparing action, leading to some distortion of the offset P2 component (as seen in Figure 4), making the comparison of P2 across durations less ideal. Given this, to separate the decision-making process from temporal encoding in a bisection task, we added a 300 ms gap before prompting a decision in Experiment 2. In addition, to generalize the contextual modulation, we applied ensemble context instead of sample spacing.

## Experiment 2

## Method

### Participants

20 participants with no hearing impairment took part in Experiment 2 in return for a monetary incentive or course credit at LMU Munich. The sample size was the same as in Experiment 1. All participants were naive to the purpose of the study and gave written informed consent before the formal experiment. The study was approved by the Ethics Board of the Department of Psychology at LMU Munich.

Because of the excessive eye or body movement artifacts during EEG recording, three participants were excluded from further analyses. Thus, the results of 17 participants (6 females, mean age 27.3 years, *SD* = 3.5 years) were reported here.

### Stimuli and Procedure

The experimental setup was the same as in Experiment 1, with the following two exceptions: first, a 300-ms blank was inserted between the stimulus offset and the question mark (prompting for a response), providing a decision time buffer for short durations (Fig. 5a); second, two sessions had the same equal-spaced duration set of [400, 600, 800, 1000, 1200, 1400, 1600] ms, but sampled with different frequencies (Fig. 5b). In one session, the above durations were tested [12, 24, 36, 48, 60, 72, 84] times, respectively. We referred to this session as the ascending frequency (AF) session. In the other descending frequency (DF) session, the same durations were tested [84, 72, 60, 48, 36, 24, 12] times, respectively. Within each session, the durations were randomly selected with the respective frequency. The order of sessions was counterbalanced among participants (before the outlier exclusion).

**Figure 5.**
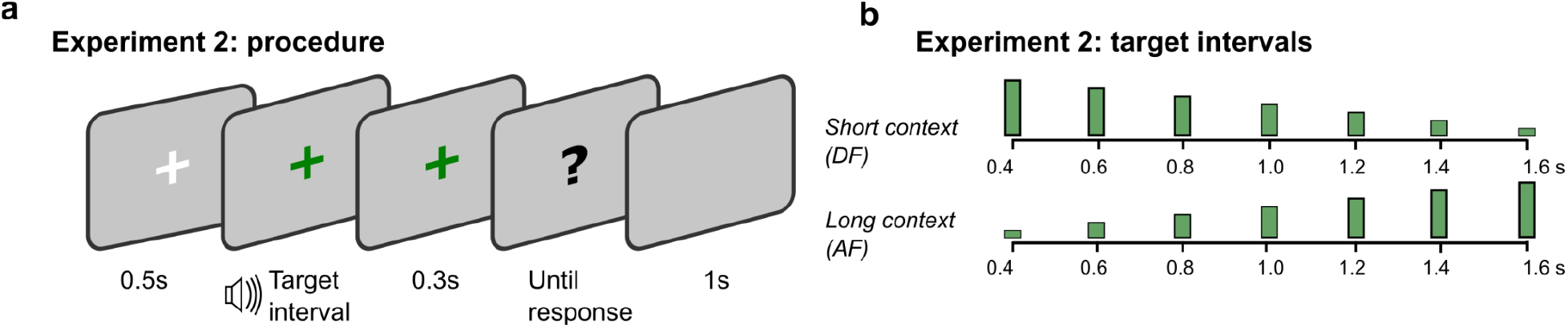
**a)** Each trial started with a fixation cross for 500 ms, followed by a target interval presentation. 300ms after the presentation, a question mark was presented, prompting participants to respond. The inter-trial interval was 1000 ms. **b)** Target intervals used in Experiment 2. In the short context session (DF), equally spaced intervals between 400 ms and 1600 ms were presented 84, 72, 60, 48, 36, 24, and 12 times during the session, whereas the presentation frequencies were mirrored in the long context session (AF).

### ERP components

In addition to the analyses of the CNV slopes, latencies, and amplitudes, we compared the CNV activities between two sessions for the intermediate target intervals (800, 1000, and 1200 ms), where the temporal context greatly modulated the bisection decision. The LPCt components were estimated on the same frontocentral electrodes as the CNV analysis (Damsma et al., 2021), but baselined relative to the 100 ms time window surrounding the onset of the question mark (50 ms preceding and following the question mark) (Kononowicz & van Rijn, 2014). We extracted the LPCt peak latencies as the maximum (most positive) amplitude within the 500 ms window starting from the question mark and calculated the LPCt mean amplitudes by averaging waveform between 300 and 500 ms after the stimulus offset (Bueno & Cravo, 2021; Ofir & Landau, 2022).

## Results

### Behavioral results

Figure 6a illustrates the averaged proportion of long responses and corresponding estimated psychometric functions. The mean PSE *(±SE)* was 749.3 ± 33.86 ms for the DF session, significantly shorter than the AF session (951.2 ±49.91 ms), *F*(1, 16) = 27.72,*p* < .001, 2 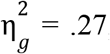, *BF* > 100, indicating the durations m the DF session were perceived longer than the *9* same duration in the AF session. Thus, the finding is consistent with the previous study (Zhu et al., 2021). Moreover, the mean JND (±*SE*) was 96.3 ± 6.69 ms for the DF session, significantly smaller than the AF session (119.8 ± 12.09 ms), *F*(1, 16) = 8.82, *p* = .01, 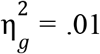, *BF* = 5.22, showing that the sensitivity of the bisection was higher in the AF compared to the DF session.

**Figure 6.**
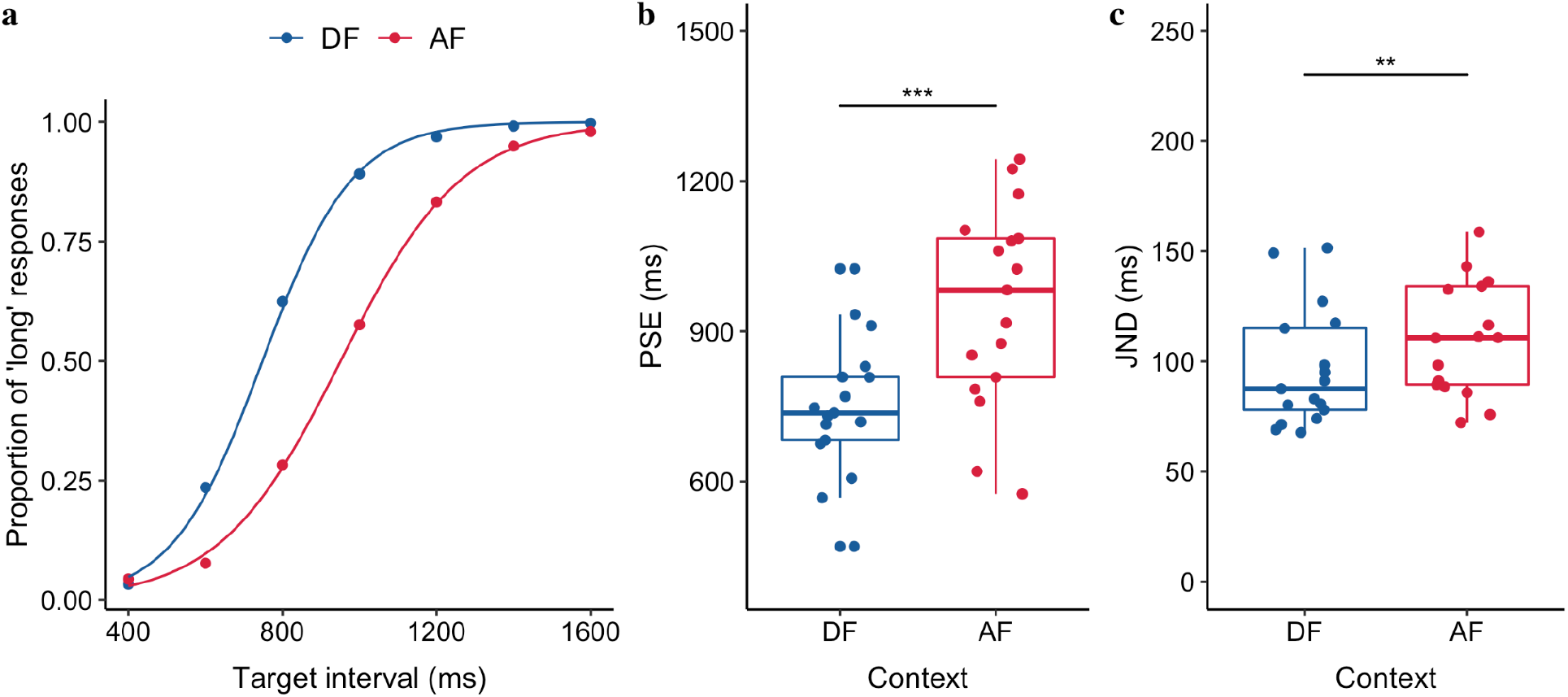
**a)** Bisection functions (proportions of “long” responses plotted against the target durations, and fitted psychometric curves) averaged across 17 participants for the two, descending (DF) and ascending frequency (AF) distributions. **b)** Boxplots of PSE of the duration judgments for the DF and AF sessions (*** p < .001). The dots depict individual PSEs estimated from individual participants. The lower and upper tips of the vertical lines correspond to the minimum and maximum values, the box represents the interquartile range (between 25% and 75%), and the horizontal line represents the median. **c)** Boxplots of JND of the duration judgments for the DF and AF sessions (** p <. 01). The dots depict individual JNDs of individual participants.

### Electrophysiological results

#### The CNV

Figure 7 illustrates the CNV activities both in the short DF (a) and long AF (d) contexts, showing that the negativity changes over time for different target intervals. Just like in Experiment 1, to characterize the CNV component, we looked into its formation rate, peak latency, amplitude, and mean latency. Given the negative ballistic deflation of the activities after P2, we used linear regression to estimate the rate (i.e., slope) at which the CNV was forming within the time window from 250 (after P2) to 650 ms (the start of the CNV). The mean values are listed in Table 1.

**Figure 7.**
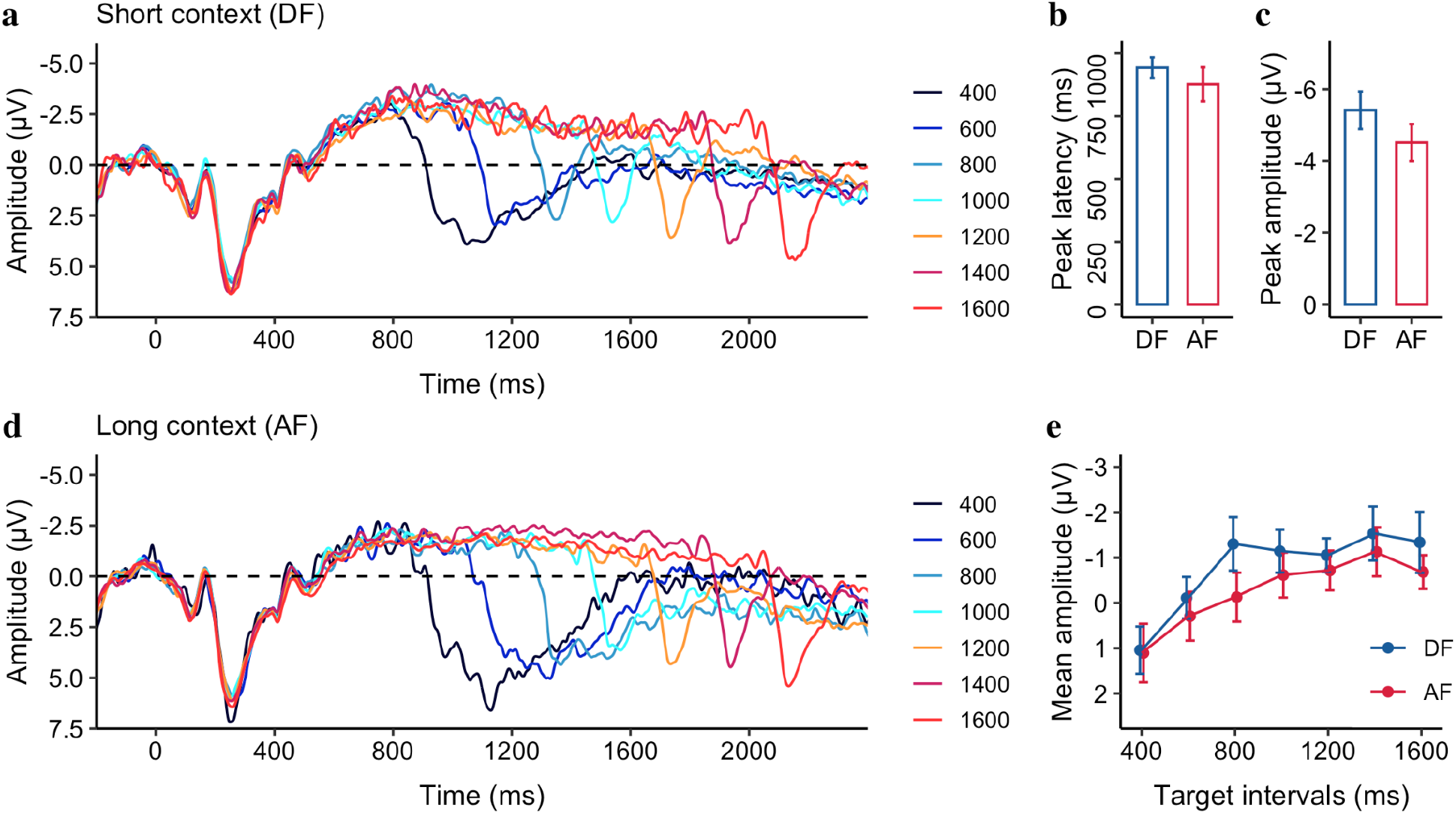
Grand average of the ERP waveforms over the medial frontal electrodes (FCz, FC1, FC2, C1, C2, and Cz) relative to the onset of stimuli, separate for **(a)** the short (DF) and **(d)** the long (AF) contexts. The mean CNV peak latency **(b)** and amplitude **(c)** of the target intervals, separated for the DF and AF conditions. (**e)** The mean CNV amplitude as a function of the target interval, separatedfor the DF and AF conditions. Error bars represent the standard error of the mean.

We found the slope was significantly negative for both the DF context [−19 ±1.5 μV/s, 95% CI = [−22, −15] μV/s, *t* (16) = −12.57, *p* < .001] and the AF context [−17 ± 1.5 μV/s, 95% CI = [−20 to −14] μV/s, *t* (16) = −11.37, *p* < .001], but significantly smaller in the DF compared to AF context, *F(1*, 16) = 5.76, *p* = .03, 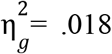, *BF =* 2.14, indicating a faster temporal accumulation in general for durations in the DF relative to the AF session, consistent with the previous research (Macar & Vidal, 2004). Moreover, the CNV peak amplitudes were significantly higher (−5.4 μV vs. −4.5 μV) for the short context (DF) relative to the long context (AF), *F*(1, 16) = 5.89, *p* = 0.03, 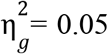, *BF* = 2.22), but with a comparable latency (942.8 ms vs. 876.6 ms, *F*(1, 16) = 0.85, *p* = 0.37, 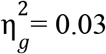, *BF* =0.49). *g*

Next, we applied a linear mixed model to the mean CNV amplitude, with the Context as the fixed effect and Duration as a covariant, which showed that the mean negativity amplitude increased by 1.66 pF for each second increased in Duration *(b* = −1.66, *CI* = [−2.35, −0.96], *p* < .001), demonstrating again that the CNV amplitude is correlated to the target interval. However, there was no significant Context *(p* = .47) or interaction between the Duration and Context *(p* = .71).

#### Late positive component of timing (LPCt)

Next, we looked into the later positivity components, such as P2 and LPCt, in the window of [0, 500] ms. Unlike Experiment 1, we failed to find any significant difference in the P2 component (the mean amplitudes were 4.2 ± 0.5 and 4.4 ± 0.6 for the DF and AF, respectively, *p* = 0.28. There was no significant difference among different target intervals, *p* = 0.99), but as seen in Figure 8, there were visible differences in the late time window. Thus we focused on the analysis of the LPCt component. The averaged LPCt peak latencies (± standard error, *SE)* were 248.6 ± 14.03 ms and 265.9 ± 11.87 ms for the DF and AF context, respectively (Fig. 8b). Same as in CNV analysis, we applied a linear mixed model to the LPCt peak latency, with the Context as the fixed effect and Duration as a covariant effect. The mixed model showed significant effects of Context (*b* = 38 ms, *CI* = [13, 63], *p* = .003), Duration (*b* = −77 ms/s, *CI* = [−133, −22], *p* = .009) and the Duration x Context interaction (*b* = −30, *CI* = [−53, −6], *p* = .013). The LPCt peaked earlier for the short DF than the long AF context, and the latency decreased as the duration increased (Table 1). The interaction was likely owing to the comparable peak latencies between the two contexts for the long but not for the short durations (see Figure 8b).

**Figure 8.**
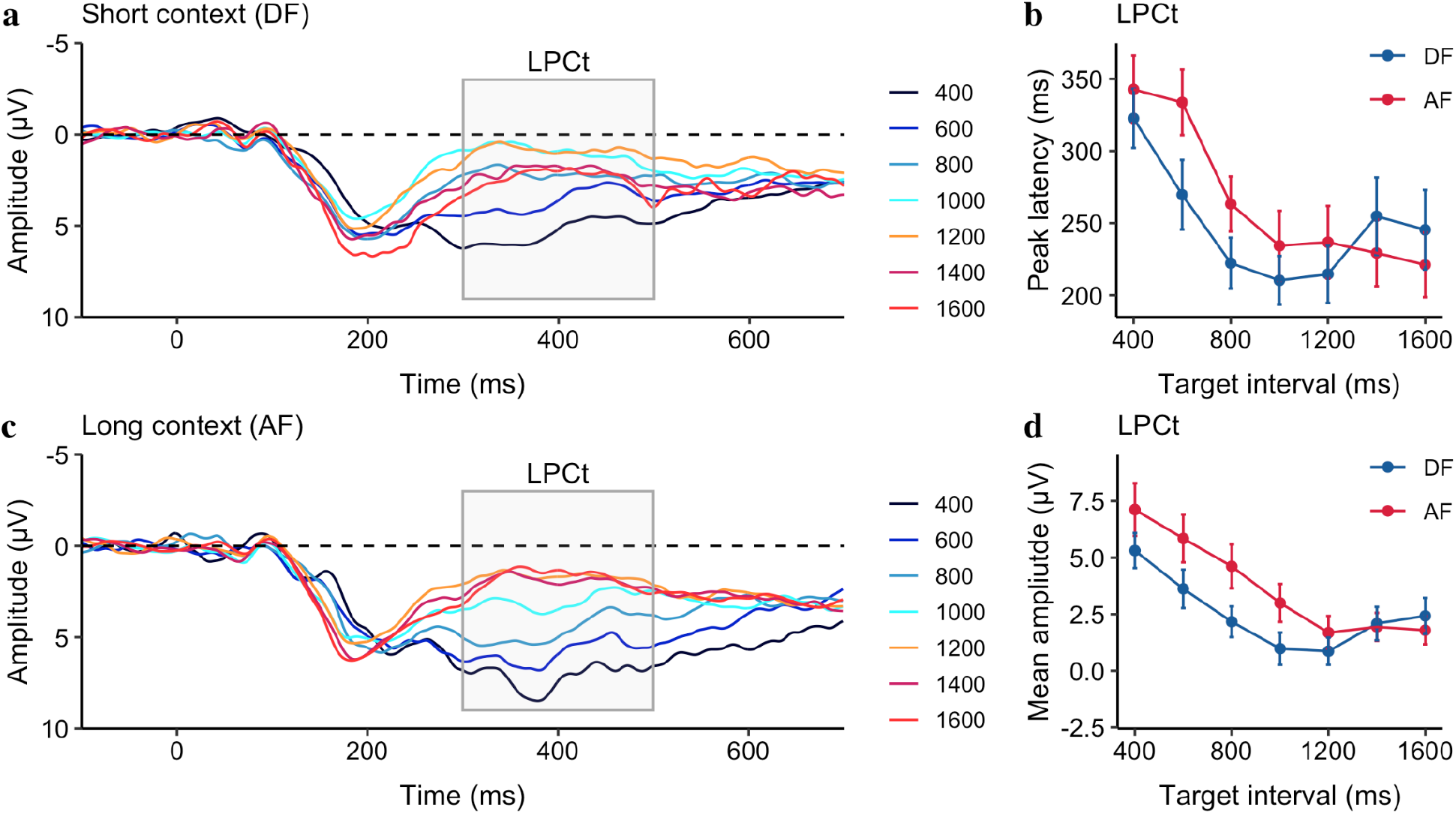
Grand average of the ERP waveforms over the medial frontal electrodes (FCz, FC1, FC2, C1, C2, and Cz) relative to the stimulus offset in the DF **(a)** and the AF conditions **(c). b)** Mean LPCt peak latency and **(d)** mean LPCt amplitude as a function of the target interval, separatedfor the DF (blue) and AF (red) contexts. Error bars represent the standard error of the correspondent mean.

For better comparison with the literature (Bueno & Cravo, 2021; Ofir & Landau, 2022), we extracted the mean LPCt amplitude from the time window of [300, 500] ms. A similar linear mixed model on the LPCt mean amplitude (see Fig. 8d) revealed similar results: significant effects of Duration *(b* = −3.55, *CI* = [−5.13, −1.97], *p* < .001), Context *(b* = 1.83, *CI* = [1.09, 2.58], *p* < .001) and the Duration × Context interaction (*b* = −1.22, *CI* = [−1.92, −0.53],*p* < .001). The mean amplitude was larger for the long AF than for the short DF context (Table 1). As seen in Figure 8d, the interaction was caused by different amplitudes for the short durations but plateaued at a similar level for the long durations.

### Cross-experiment comparisons

To gain a better understanding of the temporal encoding process reflected in the CNV and the decision-making process reflected in the offset P2 and LPCt, we further compared the results of our two experiments for short (400 ms), intermediate (around 1000 ms), and long (1600 ms) durations (as shown in Figure 9 a, b, and c). Visual inspection shows that the CNV peaked earlier in Experiment 1 compared to Experiment 2. More interestingly, even when the duration was the same, the offset late positivity was delayed by about 300 ms, suggesting that the late positive component is not solely dependent on the offset of the duration, but also on the onset of the response (the onset of the question mark that prompts for response). Moreover, the late positivity component did not fully emerge for the short duration (400 ms) in Experiment 1, largely owing to the disruption of the ongoing CNV with immediate prompting for a response.

**Figure 9.**
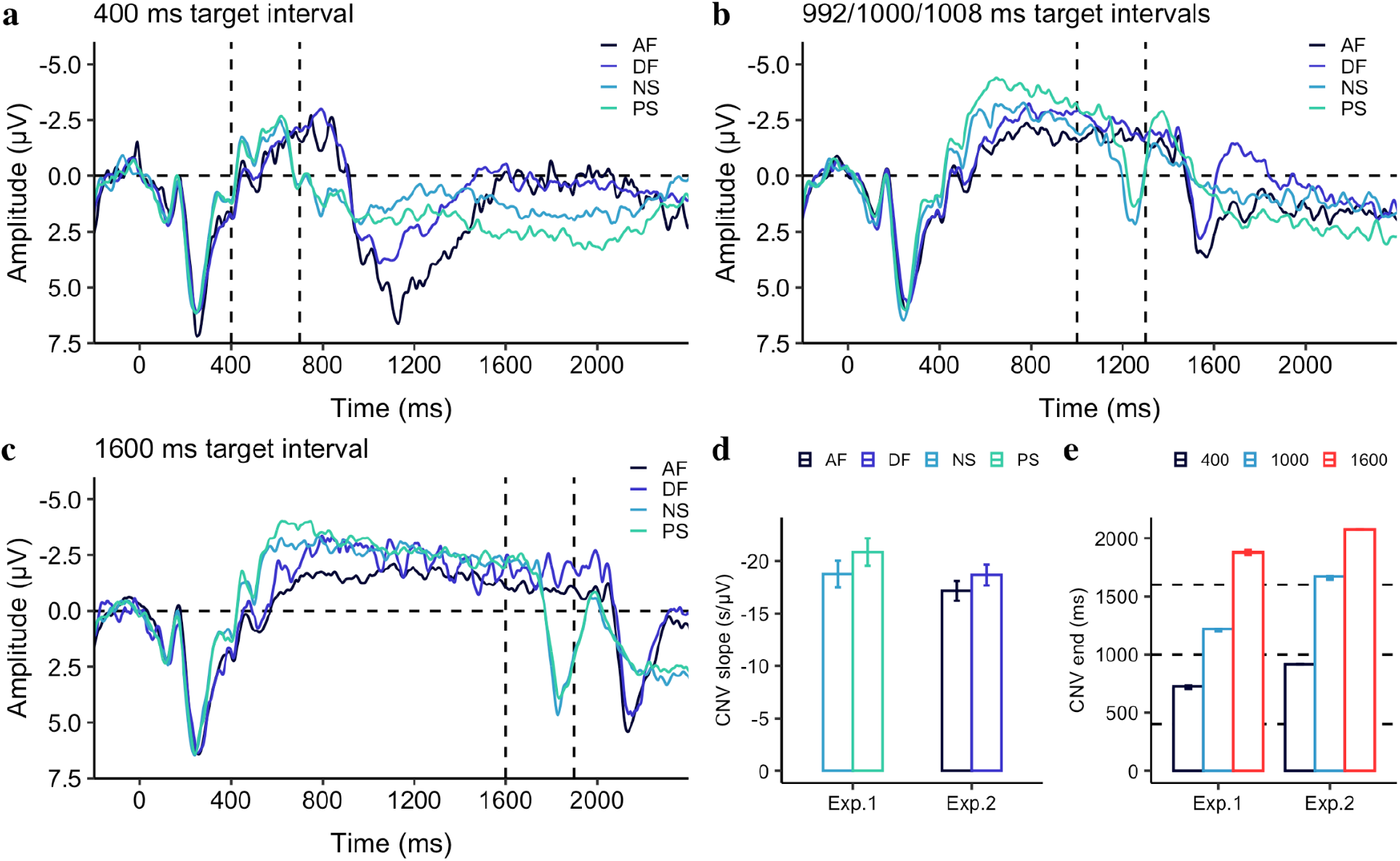
The grand average of the ERP waveforms over the medial frontal electrodes (FCz, FC1, FC2, C1, C2, and Cz) relative to the stimulus onset are depicted for the shortest **(a)**, intermediate **(b)**, and the longest **(c)** target intervals for the temporal contexts used in Experiment 1 and 2. Light blue and light green lines depict the long (NS) and short (PS) contexts of Experiment 1. Black and dark blue lines depict the long (AF) and short (DF) contexts of Experiment 2. The first vertical dashed line marks the offset cue (the question mark presentation) in Experiment 1, while the second vertical dashed line marks the offset cue in Experiment 2. **d)** The CNV slope, measured in the interval from 250 ms to 650 ms after stimulus onset. Error bars show the standard error of the mean. **e)** The CNV end, measured by the crossing point of the negativity waveform from negative to positive. Black and red colors depict the shortest and longest intervals (400 and 1600 ms) in both experiments, while the blue color depicts the intermediate durations of 992, 1000, and 1008 ms used in the experiments. For ease of visualization, they were depicted by the same color and label (1000 ms). Error bars show the standard error of the mean.

The mean slope of the CNV, measured within the interval from 250 to 650 ms, collapsed for these three intervals, was −22 ±1.1 μV/s for the short PS and −19 ± 1.0 μV/s for the long NS in Experiment 1, while in Experiment 2, the mean slope was −19 ± 1.0 μV/s for the short DF and −17 ±1.0 μV/s for the long AF. A linear mixed model was used to analyze the CNV slopes, with Experiment as the fixed effect and Context as the covariant effect. A linear mixed model was used to analyze the CNV slopes, with Experiment as the fixed effect and Context as the covariant effect. The analysis showed that there was a significant effect of Context *(b* = 1.14, *CI* = [0.04, 2.24], *p* = .042), and Experiment (*b* = 2.60, *CI* = [0.92, 4.28], *p* = .003), while there was no interaction between Experiment and Context *(p* = 0.64).

To determine when the offset of the CNV and the onset of late positivity begin, we examined the crossing latency, which is the point at which the CNV waveform changes from negative to positive after 650 ms from the onset. We found significant differences in crossing latency between the two experiments, despite using the same probe duration (all*p*s < .001): 703 vs. 914 ms for the 400-ms target interval, 1210 vs. 1628 ms for the 1000-ms target interval, and 1860 vs. 2074 ms for the 1600-ms target interval for Experiments 1 and 2, respectively. This suggests that the CNV is not solely based on the target duration, but also reflects the expectancy of the temporal period before the decision.

## Discussion

Similar to the results of Experiment 1, we found that the mean CNV amplitude increased as the target interval increased. A comparison with Experiment 1, however, revealed the CNV is not solely based on the target interval, but also depends on the period before the decision is prompted. Interestingly, we found the rate of the CNV formation and the peak amplitude of the CNV were dependent on the context. Combining the analysis of Experiments 1 and 2 revealed that the rate of the CNV is a robust indicator of context modulation. Specifically, the short context resulted in a faster rate of CNV formation, meaning the CNV began earlier in the short context compared to the long context. Our results are consistent with the notion that CNV activity reflects not only the temporal accumulator of an internal timing mechanism (Macar et al., 1999) but also temporal anticipation (Elbert et al., 1991; Ng et al., 2011), particularly for the forthcoming decision-making (Kononowicz & Penney, 2016).

Interestingly, the offset late positivity component was more distinct in Experiment 2 than in Experiment 1, even for the short durations such as 400 ms, when we inserted a 300 ms gap before prompting for action. This suggests that the late positive component may better reflect the decision process when the CNV is fully evolved. As seen in LPCt, the mean amplitude negatively correlated with the target interval, similar to the recent findings of Ofir and Landau (2022), who reported that the offset response amplitude decreases as the interval increases, but levels off after the interval passes the bisection point. The late offset positivity components such as LPCt, P300, or P3 have been suggested as indicators of decision-making at post-perceptual stages (Baykan & Shi, 2022; Kelly & O’Connell, 2013; Ofir & Landau, 2022; Polich & Kok, 1995). For a bisection task, the decision could be made before the stimulus offset when the interval presentation passes the bisection point, as the uncertainty of the response (‘long’) is greatly reduced for long intervals compared to short intervals. For the short intervals, the online monitoring of the passage of time and comparison to the bisection point remain active (Lindbergh & Kieffaber, 2013). Thus, as suggested by Ofir and Landau (2022), the amplitude of the late positivity may reflect the distance from the decision threshold.

Most importantly, we observed contextual modulation of the late positivity LPCt component: higher amplitude but later latency for the long AF than the short DF context. Given that LPCt amplitude negatively correlated with the target duration, a higher amplitude in the long context (AF) indicates that intervals were perceived shorter as compared to the same duration in the short context (DF), closely reflecting the behavioral results.

## General Discussion

The aim of this study was to examine the timing-related ERP components to gain a deeper understanding of neural mechanisms underlying the impact of ensemble contexts on temporal judgments. Results showed that ensemble contexts, both temporal spacing and sample distribution, modulated perceived time intervals, consistent with previous studies (Penney et al., 2014; Wearden & Ferrara, 1995; Zhu et al., 2021). The PSE was biased towards the mean of the ensemble distribution, with short contexts lowering the PSE, resulting in more likely to respond “long” to the target intervals in short relative to long contexts. EEG analysis also revealed context effects on the slope of contingent negative variation (CNV) and the latency and amplitude of the late post-offset positivity related to timing (LPCt), which are commonly associated with expectancy and decision processes of timing.

### The CNV

In both experiments, we saw sustained negativity, known as CNV, emerging after the onset P2, peaking between 600–800 ms, and dissolving at the end of the stimulus presentation. The CNV has been considered a strong signal for temporal processing, and early studies have suggested that its evolving slope and amplitude reflect the passage of time (Macar & Besson, 1985; Macar & Vitton, 1982). Our results showed that long durations elicited longer sustained negativities compared to short durations. However, the CNV represents more than just timing. For example, comparing brain activity between two experiments, the sustained negativity elicited by the same duration was nearly 300 ms longer in Experiment 2 than in Experiment 1, due to the 300-ms blank period before the cue display for response in Experiment 2. This modulation of response delay by the cue display supports the early proposal that the dissolving of the CNV may also indicate readiness to act quickly (Loveless & Sanford, 1974; Naatanen, 1970).

Kononowicz and Penney (2016) have recently echoed this idea that the CNV is not just about timing, but also influenced by more complex cognitive processes, such as anticipation and expectation, as well as response preparation (Kononowicz et al., 2018; Kononowicz & Penney, 2016; van Rijn et al., 2011). For example, in a study where participants were cued to respond as quickly (speed trial) or accurately (accuracy trials), the CNV amplitude was more negative in speed trials than in accuracy trials (Boehm et al., 2014), suggesting that CNV amplitude may reflect changes in participants’ response caution favoring quick decision making. Similarly, Ng et al. (2011) showed that CNV activity for the current long interval leveled off after passing a memorized internal criterion (around the geometric mean of sample intervals). In both experiments, our results also showed that the mean CNV amplitude increased with increasing target interval, leveling off around the middle intervals (Figures 3 and 7), suggesting that the CNV amplitude is closely tied to the expected decision criterion.

Another key finding is the contextual modulation of the climbing rate of CNV. In both experiments, the short context led to faster CNV formation compared to the long context. Since the CNV rate was already determined at the beginning of the presentation when the length of the stimulus was unknown, it reflects the general expectation of how long the decision interval (the ensemble mean of the sample distribution) may arrive in a given block. Thus, the rate difference between short and long contexts indicates whether the internal decision interval shifts earlier or later. While we observed context differences in peak latency in Experiment 1 and peak amplitude in Experiment 2, as well as some numerical differences in the mean CNV amplitude, the effects were not consistently significant across both experiments.

Together, our results suggest that CNV reflects the readiness or expectation to respond to an incoming stimulus (Boehm et al., 2014; Kononowicz & Penney, 2016; Ng et al., 2011), and the rate of initial CNV formation is a good indicator of context modulation of the decision threshold.

### The offset positivity components (P2 and LPCt)

After prompting for a response, we saw an offset positivity waveform, peaking at 200–400 ms and lasting for over 600 ms after the stimulus presentation. This offset positivity is known as P2 (Kononowicz & van Rijn, 2014; Tarantino et al., 2010), P3/P3b (Ofir & Landau, 2022), or LPCt (Paul et al., 2011; Wiener & Thompson, 2015) depending on studies. Depending on the timing of the response cue, either immediately after the duration stimulus or after a 300-ms gap, we observed an offset P2 (no gap) or LPCt (with a gap) that were related to the temporal decision. Short intervals, relative to long intervals, elicited delayed latency for both P2 and LPCt, and higher amplitudes for LPCt.

The early findings of time-related offset P2 came from the duration comparison studies that compared a probe interval either shorter or longer than the standard interval (Kononowicz & van Rijn, 2014; Tarantino et al., 2010) - shorter intervals elicited higher amplitudes and long latencies. Using the bisection task, we only found the latency dependent on duration in Experiment 1. When a decision was requested immediately after the duration presentation, the P2 amplitude was likely influenced by ongoing CNV activity for the short intervals (e.g., 400 and 504 ms). As the between-experiment comparison showed that when the decision response was delayed for 300 ms (Experiment 2), the late positivity was better evolved. However, we did not find any duration-related modulation in P2. Instead, the late positivity component LPCt had a strong relationship with test durations in the decision-making stage.

Late positive components, such as LPCt or P3/P3b in prior research have been measured relative to the response (Bannier et al., 2019; Wiener & Thompson, 2015) or the stimulus offset (Gontier et al., 2009; Tarantino et al., 2010) with prefrontal (Gontier et al., 2008; Paul et al., 2011) or centroparietal electrode sites (Bannier et al., 2019). The late post-positivity has been linked to the involvement of post-perceptual processes (Lindbergh & Kieffaber, 2013), similar to the idea that task difficulty is involved in decision processes (Gontier et al., 2009; Paul et al., 2011). For long durations, memory and decision-making processes would be already finished at the offset, whereas for short durations, these processes would still be ongoing. This means that compared to long durations, short durations resulted in higher LPCt amplitudes and longer latencies. In this study, LPCt was measured over prefrontal electrodes relative to the onset of the response cue, 300 ms after the test duration offset. The results showed that LPCt amplitude and peak latency decreased as the target interval increased and leveled off around intermediate durations, a pattern similar to a recent study (Ofir & Landau, 2022), which found that the amplitude of the late positivity correlated with the distance to the decision boundary in a drift-diffusion model (DDM). According to the DDM, the uncertainty of temporal bisection depends on the distance between the accumulated time to the decision boundary - bisection threshold. Short intervals with large uncertainty elicited high LPCt amplitudes, while long intervals with less uncertainty resulted in low amplitudes. Our findings are thus consistent with this interpretation. More interestingly, LPCt was found to be context-dependent, with short contexts leading to earlier peak latencies and lower amplitudes compared to long contexts, indicating that the decision boundary was set lower for the short context and, thus, the distance to the boundary was generally shorter.

### Context-dependent modulation

Both CNV and LPCt signals have been shown to depend on contextual modulation. Climbing of CNV activity was faster, and the amplitude and latency of LPCt were lower for short contexts compared to long contexts. Previous studies have shown temporal context can impact CNV in different ways (Damsma et al., 2021; Wiener & Thompson, 2015). For instance, in a reproduction task, Wiener and Thompson (2015) found that the CNV amplitude of a current trial was linearly shifted by the duration of the previous interval, with larger negative amplitudes for longer prior durations. In Experiment 1, we also found that the long context (NS) induced larger CNV amplitudes compared to the short context (PS). However, this was not the case in Experiment 2, where the short context (DF) elicited numerically higher amplitude than the long context (AF). This suggests that the CNV amplitude is more sensitive to short-term (e.g., inter-trial duration changes) rather than long-term (e.g., session-wise changes) context modulation. In contrast to the CNV amplitude, the rate of CNV formation was faster for short contexts compared to long contexts in both experiments. The CNV and climbing neuronal activity are believed to have a close relationship (Pfeuty et al., 2005), and the formation of CNV indicates how the brain encodes the timing of an upcoming event. In this study, the rate of CNV reflected the expectation of the decision threshold, which was influenced by the ensemble context.

The climbing CNV activity develops early in the perceptual encoding stage, which is tied to the memory representation of the internal criterion. In contrast, the formation of LPCt occurs during the decision stage, reflecting the comparison process of the perceived duration and the internal criterion. In this study, we showed that context affects the uncertainty of the comparison by altering the PSE towards the ensemble mean. This reduces the uncertainty of bisection for the short context in general as the test duration reaches the threshold earlier in the short relative to the long context. As a result, the amplitude and latency of the LPCt decrease. It is worth noting that the context-dependence of the amplitude and latency of the LPCt has been documented in previous research. For example, Ofir and Landau (2022) found that the late positivity remains similar in both short-range (subsecond) and long-range (supra-second) bisection tasks, even though the duration considered “short” in the long-range is longer than all durations in the short-range.

## Conclusion

In this study, we found that ensemble context, both sample spacing and frequency, impacted the bisection task, shifting the bisection point towards the ensemble mean. Temporal context modulation was also evident in the changes in ERPs related to interval timing. In the short context, compared to the long context, the CNV climbing rate increased, and the amplitude and latency of the LPCt were reduced. Both CNV and LPCt were linked to the given test duration, but were not limited to absolute durations. Our findings, consistent with the previous studies (Baykan & Shi, 2022; Boehm et al., 2014; Ofir & Landau, 2022), indicate that the CNV represents an expectancy wave for upcoming decision-making, while LPCt reflects the decision-making process, both CNV and LPCt influenced by the temporal context.

## Data availability

The data supporting the findings of this study and the code of the statistical analysis used in the manuscript are available at G-Node (DOI: 10.12751/g-node.7snfwg).

